# The monologue of the double: allocentric reduplication of the own voice alters bodily self perception

**DOI:** 10.1101/2020.08.11.246397

**Authors:** Marte Roel Lesur, Elena Bolt, Bigna Lenggenhager

**Author notes:** Corresponding authors: Marte Roel Lesur, Binzmühlestrasse 14, Box 9, Zurich, 8050, Switzerland. and Bigna Lenggenhager, Binzmühlestrasse 14, Box 9, Zurich, 8050, Switzerland.

## Abstract

During autoscopic phenomena, people perceive a double of themselves in extrapersonal space. Such clinical allocentric self-experiences often co-occur with auditory hallucinations, yet experimental setups to induce similar illusions in healthy participants have been limited to visual doubles. We investigated whether feeling the presence of an auditory double could be provoked in healthy participants and how it might affect spatial aspects of bodily self-consciousness. We recorded the own versus another person’s voice while walking around in the room using binaural headphones from an egocentric and an allocentric perspective. In comparison to listening to their own moving voice egocentrically, when listening to themselves allocentrically, participants reported a strong feeling of a presence with a similarly high degree of self-identification, suggesting a successful induction of the feeling of an acoustic doppelganger. When pointing to the source of the own voice participants localized it closer to themselves than when listening to another person’s voice, suggesting a change in spatial perception. Interestingly, the opposite pattern was found in participants that had previous hallucinatory experiences. These findings show that listening to one’s own voice allocentrically can manipulate bodily self-consciousness and self-related spatial perception. This paradigm enables the experimental study of the relationship between auditory vocal hallucinations and bodily self-consciousness, bridging important clinical phenomena and experimental knowledge.

## Introduction

The subjective location and spatial boundaries of the body are considered fundamental aspects of bodily self-consciousness (Blanke et al., 2015). Yet, they might be dramatically altered in clinical conditions. During autoscopic or doppelganger phenomena, for example, people perceive themselves or a reduplication of themselves in extracorporeal space. These experiences are accompanied by a strong self-identification with the double and bodily sensations (Brugger et al., 1997). In an effort to better understand such conditions and underlying neurophysiological mechanisms, experimental setups to induce similar states of altered bodily self-consciousness in healthy individuals have been developed (Ehrsson, 2007; Lenggenhager et al., 2007). One such case is an allocentric full-body illusion, where participants see a body in front of their actual location being stroked on the back while they are simultaneously stroked on their own back (Lenggenhager et al., 2007). These multisensory cues stimulate self-identification with the allocentrically seen body, resulting in changes in self-location and peripersonal space (Lenggenhager et al., 2009; Noel et al., 2015). Similar paradigms have almost exclusively used visual capture of other sensations to evoke such autoscopy-like illusions, which reflects the prevailing role of vision in the experimental study of bodily self-consciousness. However, clinical evidence suggests a strong link between disorders of bodily self-consciousness and conditions with prevalent auditory vocal hallucinations (Hugdahl et al., 2008; Klaver & Dijkerman, 2016; Salomon et al., 2020), and autoscopic phenomena are often accompanied with auditory hallucinations, including auditory doppelgängers (Lukianowicz, 1958; Posey & Losch, 1983). Notably, not only psychotic individuals seem more susceptible to alterations of bodily self-consciousness (Klaver & Dijkerman, 2016; B. Nelson et al., 2014), but prevalent symptoms in the condition are both auditory vocal hallucinations (Alderson-Day et al., 2020) and abnormal bodily self experiences (Di Cosmo et al., 2018; Nelson et al., 2013). These results point to a link between altered self-consciousness and self-related auditory-vocal processes, which have, however, been seldom addressed directly in the above mentioned experimental research. Here, we developed a protocol to systematically investigate the vocal and acoustic contribution to alterations of bodily self-consciousness by duplicating healthy participants’ acoustic self (voice, movement, and footsteps) in allocentric space. Blindfolded participants either heard themselves (self-allocentric) or another gender-matched person (other-allocentric) in extracorporeal space, or heard themselves egocentrically (self-egocentric). We assessed explicit and implicit changes of their sense of body and feeling of a presence, as well as implicit changes in self-related spatial perception. Participants were expected to report a similarly higher feeling of a presence for both allocentric conditions compared to the egocentric one, and a similarly higher degree of self-identification for both self-conditions compared to the other-allocentric condition. We anticipated a temperature reduction for the self-allocentric condition compared to the others, as similar effects have been found for experimentally induced autoscopy (Salomon et al., 2013), allegedly due to a disruption of self-related homeostatic processes (Moseley et al., 2008; Salomon et al., 2013). Increased heartrate for the same condition was expected as a reflection of emotional arousal (Kokkinara & Slater, 2014). We further predicted a reduction in perceived distance between the participants’ location and the sound source for the self-compared to the other-allocentric condition based on previous findings from experimentally induced autoscopy, presumably following self-identification with the extracorporeal cues (Lenggenhager et al., 2007, 2009; Noel et al., 2015). It has been suggested that schizotypal traits are linked to lower self-other discriminability (Asai, 2016), and early psychotic individuals with passivity experiences show more difficulty recognizing a recording of their voice during experimentally induced feeling of a presence than those without passivity experiences (Salomon et al., 2020). Following this, a smaller difference was expected between self- and other-allocentric stimulation for participants with previous hallucinatory experiences.

## Methods

### Participants

26 right-handed healthy volunteers participated in the experimental procedure, data from one subject was removed from all the measurements due to technical problems during the audio recording, with 25 remaining (5 males; 25.3 ± 4.36 years). Participants provided informed consent and received either course credit or financial compensation. All protocols were approved by the Ethics Committee of the Faculty of Arts and Social Sciences at the University of Zurich (Approval Number 17.12.15). The studies were performed in accordance with the ethical standards of the Declaration of Helsinki.

### Experimental apparatus

Two Roland CS-10EM In-ear Monitors (Roland Coorporation, Japan) were used as binaural microphones to record the sound, and as earphones to play it back and for online monitoring. A Zoom H6 Handy Recorder (Zoom corporation, Japan) was used as an analogue-to-digital converter for the allocentric recordings, and a Zoom H1n (Zoom Coorporation, Japan) for the egocentric recordings. Both converters recorded at a rate of 48 kHz and 24 bits. Audio management, including, synchronizing, recording, monitoring and playback was performed with software developed using Max 7 (Cycling 74, CA, USA). This software was running on a Macbook pro computer (Apple, CA, USA) with 2.3 quad-core intel Core i7 processor and 16 GB of RAM running OS X 10.9.5 Mavericks. A 3D printed dummy head with a pair of gum binaural ears attached (Free Space Pro II; 3DIO, WA, USA; the ears where used without the built-in microphones) with the binaural microphones were positioned inside the ears for recording the allocentric sounds.

For visual stimulation and interacting with a virtual 3D space, an Oculus CV1 HMD (with detached headphones) was used together with an Oculus Touch controller and four Oculus Sensors (Oculus VR, Irvine, CA, USA). The sensors tracked the HMD and the controller with 6 degrees of freedom. The control system was designed using Unity 2017 for displaying the 3D virtual room, displaying a 3D pointer, mapping and storing the 3D coordinates of the Oculus Touch controller, sending triggers to the audio software, and displaying the questionnaires in a computer screen. The system ran on a PC (Nvidia Geforce GTX 1070 8GB; 16GB RAM; Intel Core(TM) i7-8700, 3.2 GHz; Windows 10). The questionnaires were displayed and answered using a computer monitor, and a mouse.

The electrocardiac signals were recorded using an Arduino Uno (Arduino AG) and a e-Health Sensor Platform 2.0 (Libelium Comunicaciones Distribuidas S.L., Spain) with a sampling rate of 62.37 Hz. Temperature was recorded with a HH309A Data Logger thermometer (Omega, Stanford, CT, USA) at a 0.5 Hz sampling rate.

### Procedure

#### Stimuli preparation

Two sets of binaural microphones were used for recording spatially accurate tridimensional sounds from different perspectives. One set was positioned in the ears of a dummy head localized in the experimental room at a height of 1.45 m, corresponding to the participants’ position during the stimulation part of the experiment. From this perspective, the participants, as well as a gender-matched (prerecorded) stranger, were recorded while speaking out loud and walking in the room (self-allocentric and other-egocentric conditions, respectively). Both were recorded following the same actions. With the other set of binaural microphones, we simultaneously recorded sounds from the ears of the walking and talking participants (self-egocentric; Figure 1a). For a first experimental block participants were recorded opening the door to the experimental room, entering and then closing the door again. Afterwards, they walked to a point marked on the floor and began reading a fragment of The Little Prince (Saint-Exupéry, 2000) while walking a path. The direction to follow for each sentence was marked on the floor and on their reading sheet. For the recording of a second block, participants were requested to repeat out loud the phrase “one, two, three, four, five; point at my feet” 18 times as they walked following a laser pointer on the floor and holding a spatially tracked virtual reality controller (Oculus Touch) on their heads. They stopped and faced the dummy head every time before saying “point at my feet”, after which they clicked a button on the controller that would track and record their position at that moment. Each trajectory was marked by the experimenter using a laser pointer.

**Figure 1.**
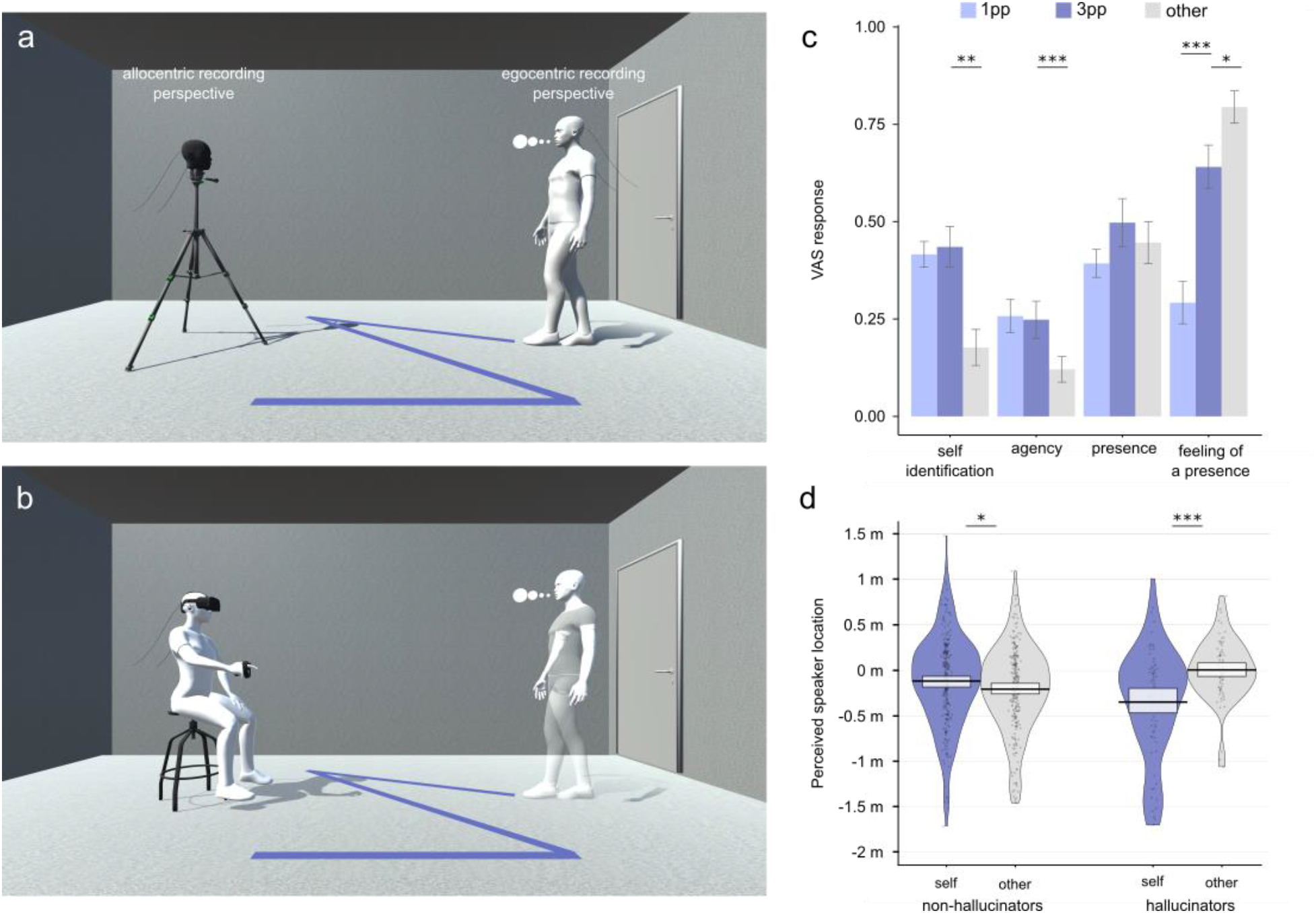
Experimental setup during the (a) recording phase, showing the perspective from which the self and other-allocentric and the self-egocentric audios were recorded. The stimulation phase (b) shows participants taking the position of the dummy head and hearing themselves or somebody else around them, at given moments they were required to point at the feet of the speaker. Plot (c) shows means and standard errors for the questionnaire dimensions, and (d) raw values, central tendencies, and distribution of the perceived distance to the speaker (self/other) separated by hallucination group.

#### Experimental block on phenomenology and physiology

During the experimental stimulation part, participants sat down with the position of their heads matching that of the dummy head during the recording using a height-adjustable stool. In the first block, they heard the recordings corresponding to the three counterbalanced conditions, each followed by a 14-item questionnaire responded on a visual analogue scale (VAS; ranging from 0 to 1, strongly agree to strongly disagree respectively) on a computer. The questionnaire was adapted from Blanke et al. (2014) and Dobricki & Rosa (2013) and included items for the *self-identification*, *spatial presence*, *agency*, and *feeling of a presence* groupings (Table 1). An additional item on voice recognition was included to confirm that participants could recognize their own recorded voice. The three conditions were included in this block to be able to disentangle the differences between self and other as well as the effect of perspective (egocentric/allocentric) between the two self-conditions. The participants’ body temperature (using two thermocouples, one placed on the back of the neck and another on the left dorsal side of the left wrist) and heart rate for each condition were recorded during stimulation.

**Table 1.** Descriptive statistics and Bonferroni corrected pairwise comparisons of individual questionnaire items, N = 23.

| Item | Self egocentric |  | Self allocentric |  | Other allocentric |  | Perspective comparisons |  |  | Self/other comparisons |  |  |
| --- | --- | --- | --- | --- | --- | --- | --- | --- | --- | --- | --- | --- |
| | Median | IQR | Median | IQR | Median | IQR | $p$ | $r$ | $Z$ | $p$ | $r$ | $Z$ |
| *Self identification* | 0.43 | 0.18 | 0.49 | 0.38 | 0.08 | 0.27 | 1.00 | -0.05 | -0.31 | 0.00 | -0.49 | -3.34 |
| Sometimes I felt like I was walking around the room myself | 0.11 | 0.29 | 0.41 | 0.57 | 0.04 | 0.17 | 0.01 | -0.40 | -2.71 | 0.00 | -0.47 | -3.16 |
| Sometimes I felt like I was in the place of the person speaking | 0.51 | 0.41 | 0.41 | 0.63 | 0.10 | 0.40 | 1.00 | -0.01 | -0.10 | 0.02 | -0.39 | -2.66 |
| Sometimes I had the feeling that the person speaking and I were one. | 0.58 | 0.22 | 0.56 | 0.40 | 0.00 | 0.19 | 0.69 | -0.14 | -0.94 | 0.00 | -0.49 | -3.34 |
| Sometimes I had the feeling to speak for myself while hearing the voice. | 0.51 | 0.42 | 0.32 | 0.46 | 0.12 | 0.33 | 0.46 | -0.18 | -1.20 | 0.01 | -0.43 | -2.92 |
| *Agency* | 0.20 | 0.27 | 0.21 | 0.37 | 0.04 | 0.15 | 1.00 | -0.05 | -0.31 | 0.00 | -0.53 | -3.60 |
| Sometimes it felt like I had control over what I said. | 0.34 | 0.44 | 0.25 | 0.57 | 0.05 | 0.18 | 0.51 | -0.17 | -1.14 | 0.01 | -0.42 | -2.83 |
| Sometimes it felt like I could have changed the content of the spoken words. | 0.10 | 0.34 | 0.05 | 0.28 | 0.03 | 0.08 | 1.00 | -0.01 | -0.06 | 0.00 | -0.47 | -3.18 |
| Sometimes it felt like I could have moved the speaking person's body if I wanted to. | 0.17 | 0.37 | 0.18 | 0.44 | 0.02 | 0.23 | 0.81 | -0.12 | -0.83 | 0.04 | -0.34 | -2.31 |
| *Spatial presence* | 0.42 | 0.25 | 0.53 | 0.51 | 0.46 | 0.36 | 0.27 | -0.22 | -1.50 | 0.93 | -0.11 | -0.73 |
| Sometimes I feel like I'm in the thick of things, not just listening | 0.43 | 0.33 | 0.65 | 0.69 | 0.51 | 0.46 | 0.76 | -0.13 | -0.88 | 1.00 | -0.06 | -0.42 |
| Sometimes I could not tell what I heard from what actually happened in the room. | 0.20 | 0.58 | 0.36 | 0.70 | 0.29 | 0.56 | 0.31 | -0.21 | -1.43 | 1.00 | 0.00 | 0.00 |
| Sometimes it seemed like I was actually attending the event. | 0.36 | 0.38 | 0.58 | 0.53 | 0.28 | 0.51 | 0.14 | -0.27 | -1.82 | 0.31 | -0.21 | -1.43 |
| Sometimes I felt physically present in the action | 0.51 | 0.36 | 0.60 | 0.41 | 0.66 | 0.37 | 0.31 | -0.21 | -1.41 | 0.66 | -0.14 | -0.97 |
| *Feeling of a presence* | 0.35 | 0.42 | 0.73 | 0.32 | 0.85 | 0.17 | 0.00 | -0.52 | -3.55 | 0.03 | -0.35 | -2.40 |
| Sometimes I felt that someone other than me and the experimenter was present in the room. | 0.17 | 0.57 | 0.69 | 0.74 | 0.82 | 0.23 | 0.08 | -0.30 | -2.03 | 0.02 | -0.38 | -2.56 |
| Sometimes I feel that someone is walking around me. | 0.18 | 0.47 | 0.87 | 0.22 | 0.88 | 0.27 | 0.00 | -0.58 | -3.96 | 0.80 | -0.12 | -0.84 |
| *Additional items* |  |  |  |  |  |  |  |  |  |  |  |  |
| I recognized the voice of the person speaking | 1.00 | 0.05 | 1.00 | 0.04 | 0.21 | 0.79 | 0.93 | -0.11 | -0.73 | 0.00 | -0.55 | -3.71 |

#### Experimental block on spatial perception

For the second block, we measured the perceived location of the heard voice while participants heard either themselves (self-allocentric) or another person (other-allocentric) speaking while walking towards and upon arrival at 18 different target positions. Unlike for the previous block, only these two conditions with multiple repetitions were necessary for our assessment in this block. Participants wore a head-mounted display (Oculus CV1) and held a virtual reality controller as they heard these sounds on the binaural headphones. They saw a laser pointer coming out from the controller and were required to point with it at the location of each of the heard targets as they saw the room’s dimensions plus an extra meter on each side on the head-mounted display (Figure 1b). They could only hear but not see the recorded speaker. After the procedure, a 2-item hallucination questionnaire was answered on a VAS (using the same scale as above). The questionnaire was adapted from Johns et al. (2002) and included an item related to hearing or seeing things that others could not, and another to hearing voices when there is nobody around. The overall procedure lasted approximately 60 minutes.

#### Data treatment and statistical analysis

Data were analyzed with R version 3.5.1 with alpha level set to 0.05, or 95% confidence intervals, and p-values were adjusted for multiple comparisons. Data were tested for normality and analyzed correspondingly. Generalized linear mixed models were using the lme4 package in R (Bates et al., 2015). Data from two participants were removed from the questionnaire analysis, one because the task was unexpectedly interrupted for one condition and another because the participant reported not having understood the instructions. From the temperature measurements, four participants were excluded due to missing data and from the ECG five participants were excluded due to data loss plus one due to noise in the recorded signal. Data from the perceived distance task of seven participants were lost due to system malfunctioning during data storage.

For the question groupings corresponding to the first block, the mean of the items contained in the group was computed. For the temperature measures, the room temperature was accounted for by subtracting it from the recordings of the two other thermocouples. A baseline was calculated as the average temperature of the first 8 s of recording for each thermocouple. This value was then subtracted from the rest of the recording, to represent the relative change in skin temperature across the stimulation period. For each condition, the average temperature change was computed over 96 s (104 – 8 s baseline) based on the duration of the shortest audio recording. For the ECG measures, the raw ECG signal was plotted to inspect the quality of the signal (one participant was removed due to noisy data). The instantaneous heart rate out of the R-peak from each ECG recording was extracted using R. Data was then filtered to exclude heartrate values above 200 and below 25 bpm (García Martínez et al., 2017). As with the temperature recordings, data for each participant was cut to match the duration of the shortest recording.

For block two, the perceived distance to the speaker was calculated by computing the Euclidean distance between the participants’ location (the middle of the stool where they sat down), and the coordinate pointed by participants for every trial. The actual distance to the speaker was calculated by determining the distance between the participants’ position and the recorded position for each trial. For these calculations, only two axes were considered, the horizontal and the depth axes; the vertical axis was kept constant as participants were asked to point at the floor so the pointed coordinate could be accurately measured. For assessing the drift in perceived location, the difference between perceived and actual distance to the speaker was computed for each of the 18 trials. One data point was removed because it exceeded the plausible values for the room size, suggesting a measurement error (> 2.5 m). For grouping participants according to their previous hallucinatory experiences, those that scored above 0.5 on either of the two questionnaire items were considered to be in the hallucination group. For assessing differences in the perceived distance to the speaker, multiple linear mixed models were fit in a step-wise manner to examine perceived location data based on our hypotheses (see supplementary materials).

## Results

Multiple corrected Wilcoxon comparisons showed significant differences for the questionnaire between the self-egocentric and self-allocentric conditions for the feeling of a presence (*W* = 13, *p* < 0.001, *r* = −0.52), between the self-allocentric and other-allocentric conditions for self-identification (*W* = 230, *p* < 0.01, *r* = −0.49), agency (*W* = 185, *p* < 0.001, *r* = −0.53) and feeling of a presence (*W* = 60, *p* < 0.05, *r* = −0.35) (Figure 1c). Results for the individual items of the questionnaire and additional comparisons are shown in Table 1. Friedman tests for temperature comparisons between conditions revealed no significant differences for neither the left wrist (χ^2^ (2) = 0.35, p = 0.84) nor the neck (χ^2^ (2) = 0.82, *p* = 0.66). A repeated measures ANOVA showed no significant differences in heartrate between the three conditions F(2, 32) = 0.184, p = 0.83).

Before fitting linear mixed models to the perceived location data, non-independence within participants was confirmed for the overall dataset (ICC(1) = 0.31). Multiple models were fit in a step-wise manner. The first fitted model included a fixed effect of condition (b = 0.01, 95% CI: −0.06, 0.07, *t* (628) = 0.233, *p* = 0.81), a second model included both the fixed effect of condition (b = 0.01, 95% CI: −0.06, 0.07, *t* (628) = 0.233, *p* = 0.81) and group (b = −0.01, 95% CI: : −0.34, 0.32, *t* (16) = −0.05, *p* = 0.96), no differences were found in terms of predicted variance between these models (*χ2*(2) = 0.03, *p* = 0.95), however a final model including both the fixed effect of condition and group showed a significantly higher predicted variance (*χ2*(2) = 32.53, *p* < 0.001). The model showed a main effect of condition (b = −0.09, 95% CI: −0.16, −0.02, *t* (627) = −2.51, *p* = 0.01) and a two-way interaction effect of condition and group (b = 0.44, 95% CI: 0.29, 0.59, *t* (627) = 5.77, *p* < 0.001). Post-hoc Tuckey tests with a single-step correction showed a significant difference for both groups between conditions, with non-hallucinators showing a drift of the perceived targets in the direction of their own body when listening to their own voice (*SE* = 0.04, *p* = 0.04) while hallucinators showed the opposite pattern (*SE* = 0.07, *p* < 0.001). We additionally estimated the effect size for the final model according to the method suggested by Johnson (2014) and Nakagawa & Schielzeth (2013) which results in a *convergence r*^*2*^ = 0.356. In order to further confirm these results, we separately compared for each group the effect of condition. Non-independence within participants was confirmed for the non-hallucinators (ICC(1) = 0.32) and hallucinators (ICC(1) = 0.31). Linear mixed models showed a main effect of condition for non-hallucinators (b = −0.09, 95% CI: −0.16, −0.02, *t* (488) = −2.5, *p* = 0.01; *convergence r*^2^ = 0.33), and hallucinators (b = 0.35, 95% CI: 0.22, 0.49, *t* (139) = 5.18, *p* < 0.001; *convergence r*^2^ = 0.42). Visual exploration confirmed a normal distribution of the residuals for each of the described models.

## Discussion

These results suggest that listening to one’s acoustic self in extracorporeal space can manipulate phenomenal aspects of bodily self-consciousness and self-related spatial perception. As hypothesized, healthy participants indicated a strong feeling of a presence of a speaking person walking around themselves while self-identifying with that person, thus reporting the perception of an auditory doppelganger. This provides evidence for an experimentally induced autoscopy-like experience in healthy participants stimulated acoustically. Importantly, our measure of self-identification was based on a previous principal component analysis (Dobricki & Rosa, 2013) and is different to voice recognition, including the sense of body ownership (e.g. Blanke, 2012) and attributing own-experiential qualities to the self-identified object. These findings on altered bodily self-consciousness confirm verbal reports from healthy participants during piloting:

> “So my experience was strange at the beginning, but soon it felt, it felt familiar and I immediately recognized myself. That it was my voice. And in some way it was also like listening to your own self, to your consciousness in a way. It seemed also very real, as if I was going around myself. Yes, it was impressive to hear myself in such way, as if I was around myself. As if you were around yourself.”
>
> “As if you feel in that person too, but at the same time you are in between and you are surrounded. So, I don’t know, I feel that I’m surrounded, but at the same time it’s myself. So I feel in two places simultaneously, let’s say.”

These strong phenomenal descriptions however were not in line with the physiological measures, which might indicate that the effect was not as strong as in previously described multisensory illusions, but might also be linked to accumulating evidence not replicating the link between phenomenal alterations of the bodily self and physiological measures such as temperature (de Haan et al., 2017; Roel Lesur et al., 2020). Yet, in consistency with previous finings for experimentally induced autoscopy (Lenggenhager et al., 2007, 2009; Noel et al., 2015), we found changes in the location of self-identified extracorporeal cues. As predicted, participants without previous hallucinatory experiences perceived their own allocentric voice closer to themselves compared to the voice of another person, presumably due to a change in self-location resulting from self-identification with the extracorporeal cues (Lenggenhager et al., 2009; Noel et al., 2015). We expected this effect to be smaller for participants with previous hallucinatory experiences due to potentially more self-other confusion, as it has been suggested that hearing voices might be related to alterations in self-monitoring and self-other distinction (Blakemore et al., 2000). However, they showed an effect in the opposite direction, locating their own voice further away compared to the other person’s. A reduced capacity to recognize the own voice has been shown for early psychotic patients with passivity experiences during somatosensory-motor experimentally-induced feeling of a presence (Salomon et al., 2020), and schizotypal individuals tend to show a narrower peripersonal space compared to controls (Di Cosmo et al., 2018), potentially related to distorted self-other discrimination (Asai, 2016). However, it’s not clear why they would show a drift further away from their location compared to when hearing someone else’s voice. A previous study with a visually-driven experimentally-induced autocopy found no differences between schizophrenic patients and controls in neither self-location nor subjective responses (Shaqiri et al., 2018), while our study is yet to be tested with patients, these results might suggest changes compared to controls that are specific to self-related vocal-acoustic signals. Extending this study to a clinical sample might add to the understanding of the relation between acoustic-vocal perceptions and distortions of the bodily self and self-boundaries. Importantly, participants with previous hallucinatory experiences scored relatively high in explicit recognition of their own voice (M=0.77, SD =0.32 for the egocentric, and M=0.78, SD = 0.27 for the allocentric condition), suggesting that this effect is not necessarily linked to lack of explicit self-recognition. Notably, auditory vocal hallucinations are often linked to a feeling of a presence (Alderson-Day et al., 2020), accompanied by bodily alterations (Alderson-Day et al., 2020; Woods et al., 2015), and appear to come from extracorporeal space (Copolov, 2004; McCarthy-Jones et al., 2014; Woods et al., 2015). These bodily and spatial aspects during auditory hallucinations are linked to primary characteristics of the bodily self. While audition has been relatively neglected in accounts of autoscopy and out of body experiences, acoustic phenomena are frequent and reports of auditory doubles exist even in non-clinical populations (Lukianowicz, 1958; Posey & Losch, 1983). The described paradigm offers new ways to manipulate and study the relation between auditory vocal hallucinations and bodily self-consciousness both in health and disease, potentially serving as a bridge between clinical and experimental knowledge in the field.

## Acknowledgements

The authors thank Martin Fröhlich and the Institute for Computer Music and Sound Technology, ZHdK for lending the binaural dummy head, Gianluca Saetta and Marieke Weijs for the statistical assessment to analyze the perceived location data and Colin Simon for recording data. The authors also thank. M.R.L and B.L were supported by the Swiss National Science Foundation (grant number: PP00P1_170511).

